# Is blue the most attractive color for bees? Exploring the attractiveness of colors in vane traps

**DOI:** 10.1101/2025.02.10.637548

**Authors:** Elizabeth Rentería, Gunnar Brehm

## Abstract

Bees play important roles in shaping ecosystems through processes like pollination. However, their populations are in decline due to habitat degradation and unsustainable agriculture. This study assesses the effectiveness of color traps for bee monitoring in two distinct yet geographically close habitats in “Jenaer Forst” natural reserve in Germany. Over a 23-week period in spring and summer 2023, we tested six trap colors (blue, yellow, white, violet, blue-yellow, and violet-white) and analyzed their performance in terms of bee abundance, diversity, and species composition. Our results indicate that violet and blue-yellow traps captured the highest bee abundance and species richness. While overall bee community composition did not differ significantly between trap colors, specific species showed color preferences. The blue-yellow trap emerged as a versatile option, potentially capturing a diverse range of species through its contrasting colors. Moreover, despite the close proximity of the two sites, differences in community composition suggest that habitat-specific factors influence bee color preferences. Minor habitat changes significantly impacted trap effectiveness, emphasizing the importance of tailored trap designs. These findings highlight the importance of considering both color contrast and habitat context when designing bee monitoring strategies and understanding bee foraging behavior. This study contributes valuable insights to improve trap designs and enhance our understanding of the interactions between bees and their environments.

## Introduction

Insects play a vital role in ecosystems, contributing to processes like nutrient cycling, predation, and pest control (Forister et al. 2019; Hartley and Jones 2008). Among these roles, pollination is crucial for many animals, including humans, for plant-based food (Forister et al. 2019). Bees, with over 20,000 species worldwide (López-Uribe 2021), are key pollinators due to their coevolution with angiosperm flowers (Rosalind and Pitts 2008).

Bees support ecosystems and agricultural productivity (Potts et al. 2016), but their population declines due to habitat degradation and unsustainable agriculture (Bates et al. 2011). In Europe, over half of the 1,965 bee species lack red list status, and up to 50% are considered nationally threatened (Nieto et al. 2014), highlighting the urgency of studying bee species for conservation strategies. Additionally, most studies focus on common species like *Apis mellifera* and *Bombus terrestris* (Chen et al. 2020; Gumbert 2000), whereas information about solitary bees and lesser economically important bees remains limited (Acharya et al. 2022). Given that different bee species may have varying foraging strategies and visual sensitivities, understanding their perception of floral signals is crucial for effective biodiversity monitoring and conservation (Abrahamczyk et al. 2010).

Bees have well-developed UV, blue, and green receptors (Chittka and Menzel 1992; Chittka 1996). While they are most sensitive to these wavelengths, their overall visual spectrum spans 350 to 650 nm. Within this range, bees cannot distinguish certain colors directly but instead rely on contrast, reflectance patterns, and saturation differences to identify flowers (Chittka and Waser 1997). This ability plays a crucial role in their foraging decisions and attraction to optimal flower sources (Acharya et al. 2022). Flowers, in response, must compete for the attention of suitable pollinators to successfully transfer pollen. This competition drives flowers to become conspicuous, offering rewards for pollinators and ensuring recognition and remembrance (Chittka et al. 1994; Weiss and Lamont 1997). For this recognition to be well-established, the association must remain constant over time (Chittka and Thomson 2001). Consequently, color preference and perception in bees appear to be a dynamic interplay of association and cognition with their food source, rather than having a strictly hardwired and unchangeable trichromatic color preference (Avarguès-Weber and Giurfa 2014).

Passive sampling methods like color traps (pan and vane traps) are simple and effective for investigating bee preferences (Abrahamczyk et al. 2010; Acharya et al. 2022). Pan traps use color to mimic flowers, while vane traps add a flight intercept function (Sircom et al. 2018), both effectively attracting diverse insect species. Color traps are often used to investigate bee preferences (Abrahamczyk et al. 2010; Acharya et al. 2022; Buffiington et al. 2021; Campbell et al. 2023; Freitas-Moreira et al. 2016; Gonzalez et al. 2020; Hall 2016). These traps are practical for long-term use across various habitats, helping determine optimal colors for the investigation of species richness and diversity (Vrdoljak and Samways 2012).

This study tests six color traps for bee monitoring in two nearby but distinct grassland habitats: While blue and yellow have commonly been used in many studies, the colors white and violet have far less often been deployed. In addition, we also examined two selected color combinations, i.e. blue-yellow and violet-white. We expected that the commonly used colors blue and yellow would attract most species and individuals, but we also expected that violet attracts at least certain bee species due to their prevalence of violet flowers. Traps with color combinations were expected to attract a mixture of species attracted to each of the colors. This study underscores the importance of diverse biodiversity monitoring methods and explores how pollinators interact with different colors.

## Materials and methods

### Study area

The study was conducted within the limits of the “Jenaer Forst” natural reserve, encompassing 541.1 ha in the central eastern region of the state of Thuringia, Germany. Situated on the outskirts of the city of Jena, this area holds the status of a protected zone within the European Natura 2000 network. The soil in the area is characterized as very stony to clay-loamy, conformed mainly by low-shell limestone (Heyn et al. 2019). The landscape is distinguished by two predominant forest types, namely, barley-beech and woodruff-beech forests. Additionally, the slope region features orchid-rich semi-dry and dry grasslands (Heyn et al. 2019).

The specific site of this study is situated in a dry-grassland (preserved by regular removal of bushes) on a rich orchid-beech forest situated on a west-facing hillside location within the reserve (Heyn et al. 2019). The investigation spanned 23 weeks from March 11 to August 20, 2023.

Two distinct habitats were selected within the reserve (Fig. 1): Haselberg (H) (50.91888° N, 11.53122° E, 369 m.a.s.l) and Haselberg-Slope (HS) (50.91831° N, 11.52873° E, 354 m.a.s.l) (all coordinates in Table S1). These habitats were only 170 m apart, but HS is located on a plateau. Both designated habitats exhibited forest clearings with grassland-type vegetation (flowering plants list from both sites in Table S2).

**Fig. 1.**
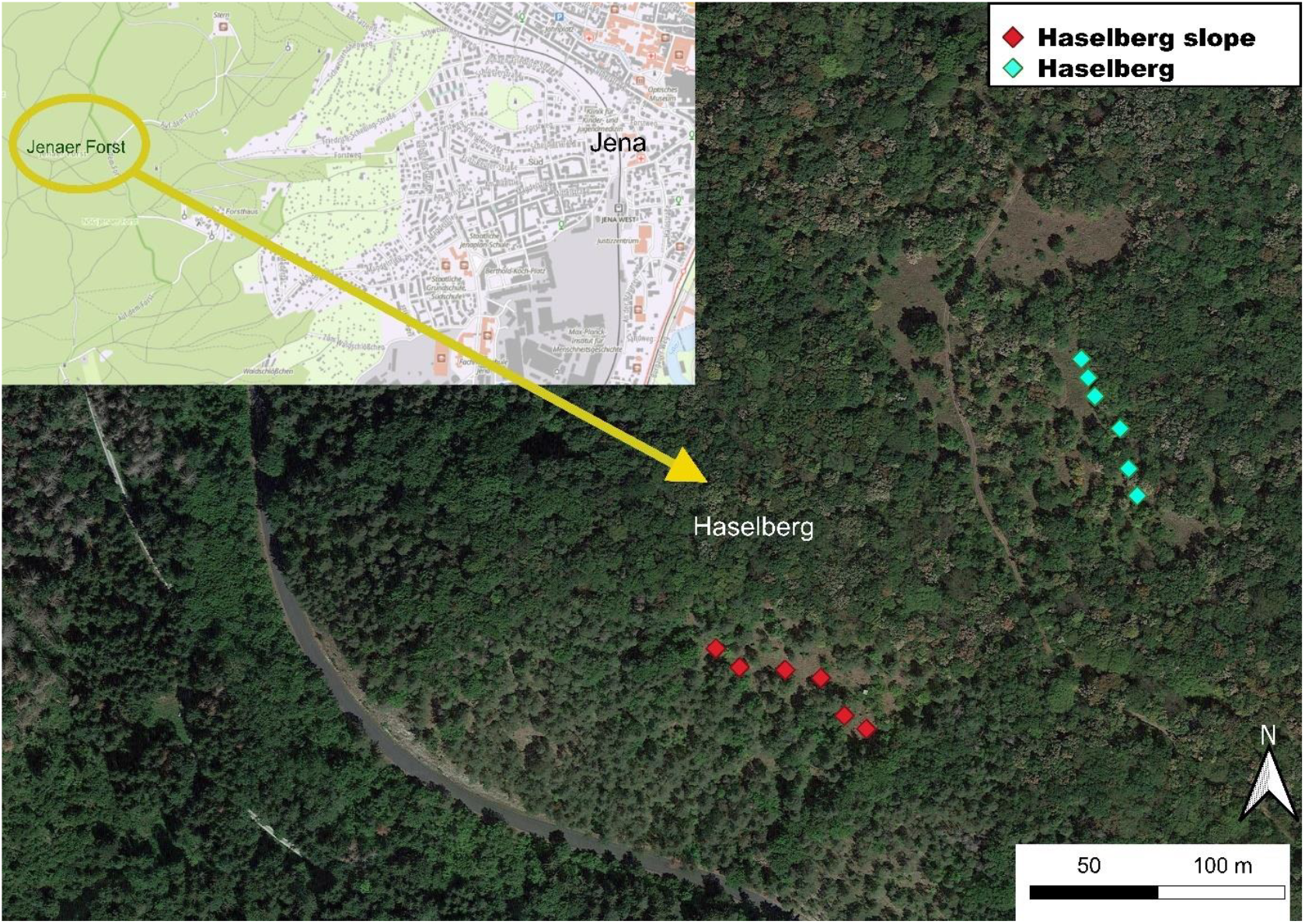
Satellite image of study area. Map created with QGIS, upper map information from © GDI - Th dl-de/by-2-0. Haselberg marked by turquoise points and Haselberg-slope marked by red points.

Environmental data was taken from the weather station of the Max-Planck-Institute for Biogeochemistry (50.909889° N, 11.571944° E), which is 3.1 km away from study area.

### Sampling

Self-constructed vane traps were used for sampling. They were designed by GB based on a modified version of light traps from Singh et al. (2022). Traps were designed as crossed vane configurations with two polypropylene panels of equal size, intersecting at the center (Fig. 2a). Each trap featured a bottom section fitted into a white funnel attached to a white bucket with lid (one liter volume), with a circular white polypropylene roof. The traps were positioned 0.7 m above the ground on a supporting aluminium structure (Fig. 2b). The traps buckets were filled with saturated saltwater to preserve captured specimens (Young et al. 2021). Saltwater was preferred over the use of ethanol because the traps were left in the field for a week, during which the ethanol would have evaporated, leading to the specimens rotting.

**Fig. 2.**
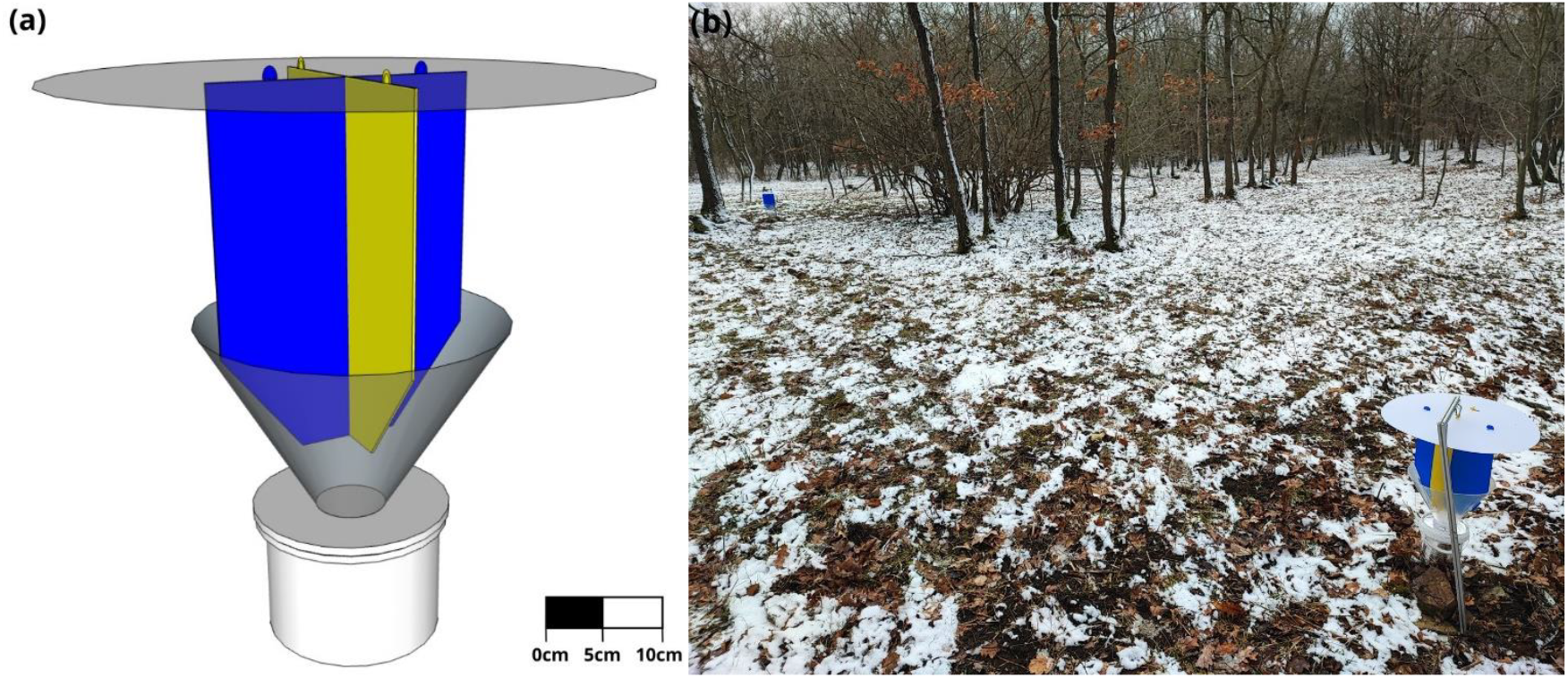
(a) Vane trap design (3D model) using the color combination blue-yellow. (b) Arrangement of traps in the study area (Haselberg) Photograph taken at the beginning of the study, early spring/March (11.03.2023). 3D model made with SketchUp.

Six variations of vane traps were made by selecting differing vane colors: blue, yellow, white, violet, blue-yellow combination, and violet-white combination. The traps were spaced at least 20 m apart, resulting in six positions in each designated area (Fig. 1). Weekly trap maintenance involved emptying and rotating traps between three of the six available positions to avoid potential bias.

Light reflectance and transmission measurements were performed by GB with a Specbos photospectrometer Blue Spec Cube (JETI, Jena, Germany). The colors reflectance was measured within a 300–1000 nm range under natural daylight conditions (Fig. S1). The specific material measured were as follows: Blue: 0.8 mm polypropylene, bright blue, from Modulor, Berlin, Germany (#3690); Yellow: 0.8 mm polypropylene yellow, from Modulor, Berlin, Germany (#4690); White: 1 mm polyoxymethylene plates; Violet: 1 mm polyoxymethylene plates, sprayed with Amsterdam spray paint, #507, ultramarine violet.

The collected bee samples were preserved in 80% ethanol. They were later prepared and curated at the Phyletic Museum in Jena, where they are permanently stored. Bees were sorted to morphospecies and identified to lowest taxa possible by JR. Further identifications were checked and made by melittologist Frank Creutzburg (JenInsekt, Jena-Germany).

### Data analyses

The statistical analysis and graphics were performed in R (version 4.3.1, 2023). First, the performance of each color variation trap was analyzed using the total number of bee specimens collected per trap, taking into consideration the position of the trap in each habitat and the collection week. The performance was analyzed using generalized linear mixed models (GLMM) with a negative binomial regression using lme4 package (version 1.1.34), followed by a Tukey’s HSD test (α = 0.05) using emmeans package (version 1.8.9). The model diagnostics to check the fit of the GLMM was performed with the DHARMa package (version 0.4.6). A Friedman test was employed to test the variation of performance of each trap variation on each position and biweekly, using the ggstatsplot (version 0.12.1) package. The time data was analyzed biweekly instead of weekly to increase legibility in the graphs.

We explored community composition through the following analysis: Initially, differences in community composition among color trap variations, habitats, and weeks were examined using a PERMANOVA. The adonis function from the VEGAN (version 2.6.4) package was employed to test for significant variations in community composition and to quantify how much each variable contributed to overall community differences. To assess the dissimilarity of the community for each variable, ANOSIM (VEGAN) was utilized, providing insight into the distinctiveness of community structures. Furthermore, to identify the significative species preferences for each trap and site, and Indicator Species Analysis (indicspecies package version 1.7.14) was employed. This method facilitated a comprehensive understanding of the factors influencing community dissimilarity. Simultaneously, community dissimilarity was visually represented through Nonmetric Multidimensional Scaling (NMDS) using the VEGAN package. The zero-adjusted Bray–Curtis coefficient, as recommended by Clarke et al. (2006), was applied to transform the data.

To understand the distribution of species within the community, an examination of species diversity was undertaken across color variation traps, habitats, and weeks. The diversity analysis involved the use of species accumulation curves (rarefaction curves) with the VEGAN (version 2.6.4) and iNEXT (version 3.3.3) packages. Through this analysis, species richness (q = 0), Shannon diversity (q = 1), and Simpson diversity (q = 2) were calculated. These three metrics were then employed to compare the richness, species evenness, and species diversity across the specified variables (Chao et al. 2014).

## Results

### Trap performance

A total of 673 bees were collected, representing 15 genera from five families and 96 species. The most abundant species were *Osmia bicolor* (Schrank,1781) (14.0%), *Bombus lucorum* (Linnaeus, 1761) (7.6%), *Apis mellifera* (Linnaeus, 1758) (7.0%), *Andrena bicolor* (Fabricius, 1775) (6.2%), and *Lasioglossum morio* (Fabricius, 1793) (4.8%). Table 1 displays the percentage of bees collected at each site and in each color trap.

**Table 1.** Percentages of bee specimens collected in each trap type.

| Site | Blue | Yellow | Violet | White | Blue-Yellow | Violet-White |
| --- | --- | --- | --- | --- | --- | --- |
| Haselberg (n=353) | 15.3 | 7.6 | **24.9** | 12.7 | 19.0 | 20.4 |
| Haselberg-slope (n=320) | 20.3 | 8.8 | 22.2 | 9.1 | **23.1** | 16.6 |
| Total (n=673) | 17.7 | 8.2 | **23.6** | 11.0 | 21.0 | 18.6 |

Analyzing the overall performance of traps for both habitats, yellow and white traps captured significantly fewer bees than most other traps (Table 1, Fig. 3). The violet trap stood out as being overall the one that collected the total highest number of bees. At the Haselberg site, only the violet and violet-white traps exhibited a significantly higher bee capture compared to the yellow trap (Fig. S3a). In contrast, at the HS site, the yellow trap was the only one that performed better than the yellow and white trap (Fig. S3b). Notably, trap performance varied between sites, despite their relatively proximity.

**Fig. 3.**
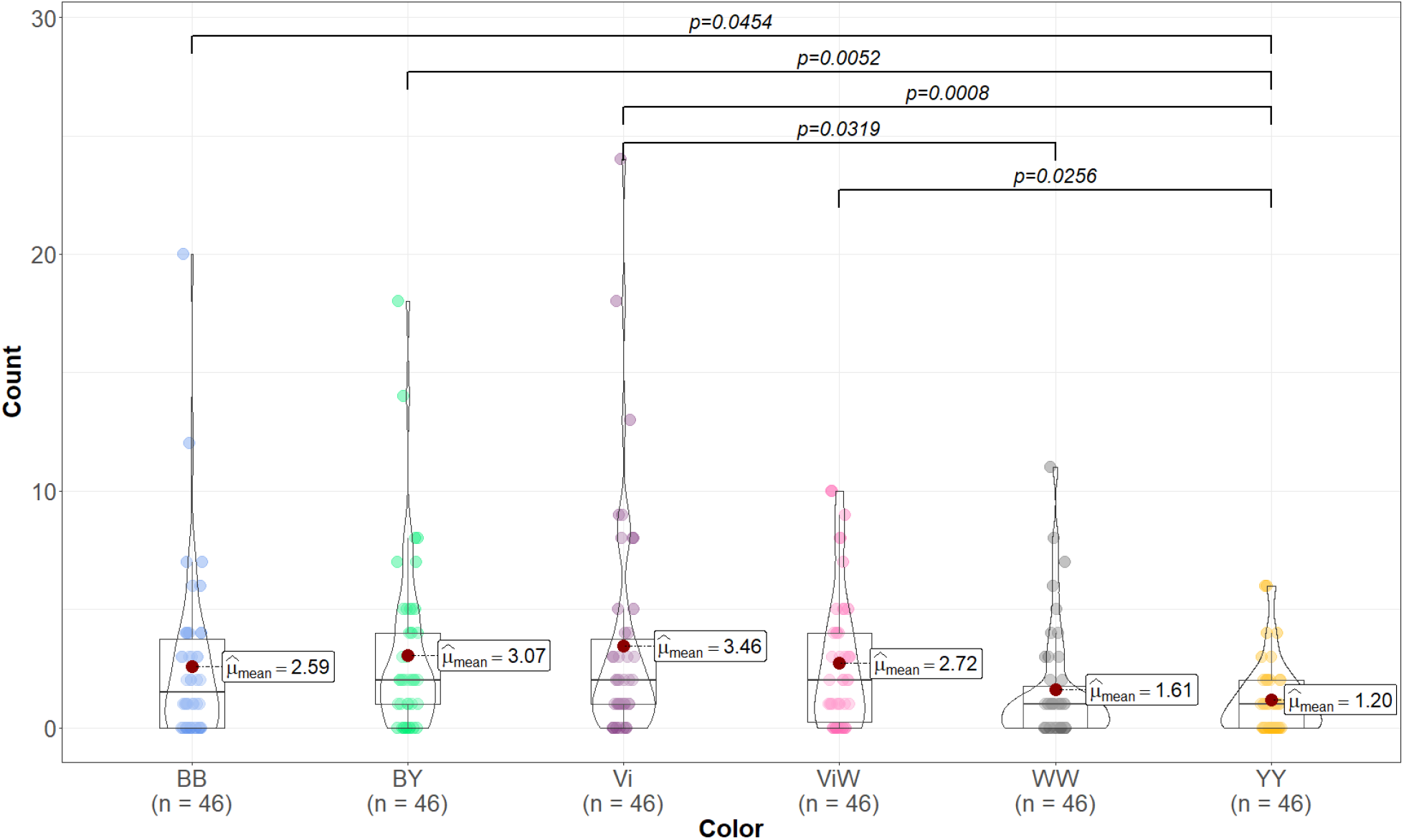
Effectiveness of color traps in capturing bees. The number of bees collected (count) by each individual vane trap is shown for both habitats (H and HS combined) over the course of 23 weeks (n=46). Each violin plot presents the mean number of the collected bees for a specific trap. Brackets indicate significant differences between traps, determined using the EMMeans pairwise comparison function with the GLMM analysis. Trap colors: BB=Blue, BY=Blue-yellow, Vi=Violet, ViW=Violet-white, WW= White, YY= Yellow.

A Friedman test was conducted to assess the performance of the traps during biweekly collection intervals. Analyzing data from both habitats combined (H and HS), substantial differences were observed in the number of bees collected, particularly between high and low collecting weeks (Fig. 4). However, no significant correlation was found between the number of collected bees and weather parameters (temperature, humidity, precipitation) during the two-week periods.

**Fig. 4.**
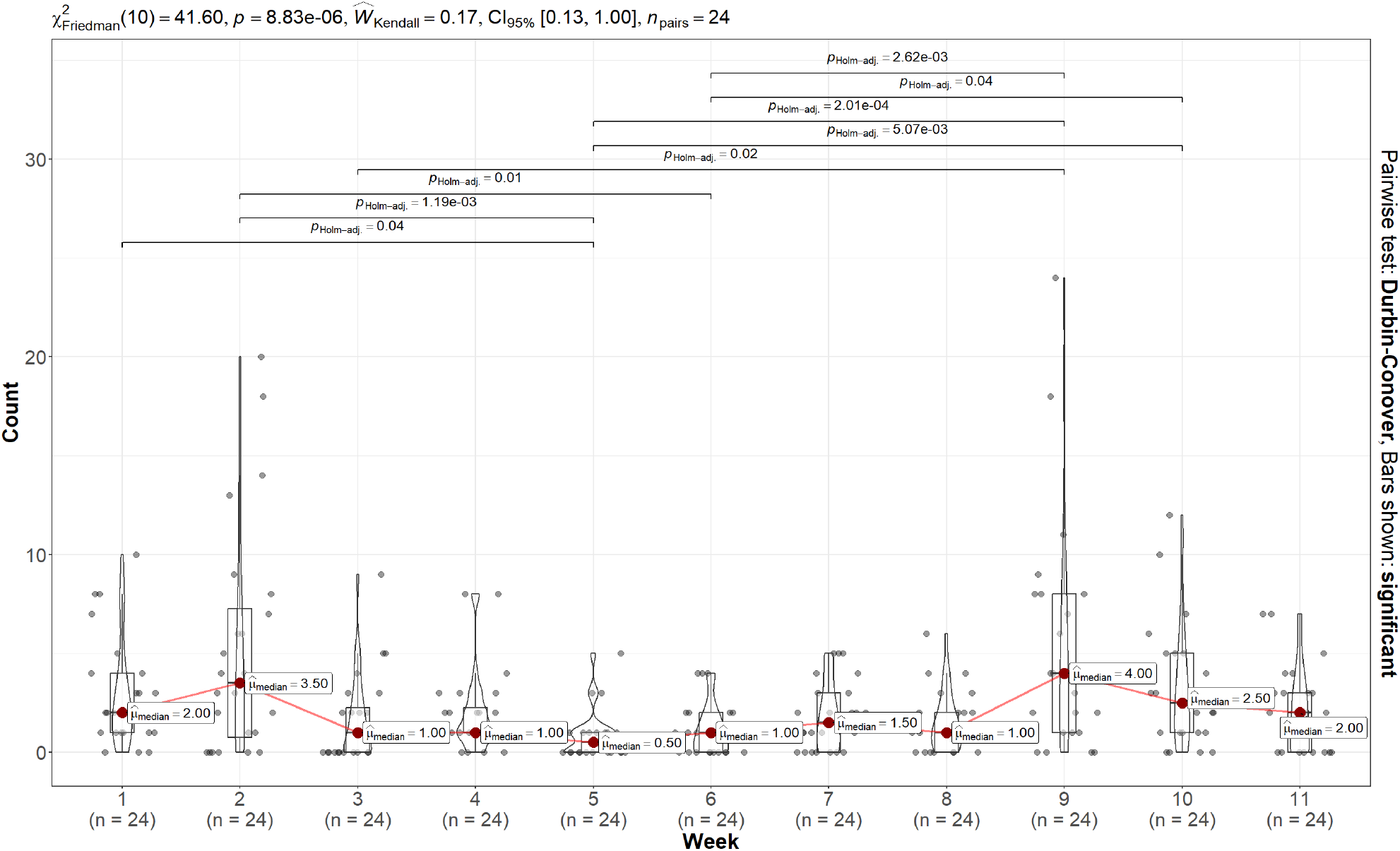
Overview of the overall performance during biweekly collecting intervals. The number of bees collected (Count) by each individual vane trap from both habitats (H and HS) is depicted for each two-week interval (n=24). Each violin plot displays the median of the collected number of bees for that specific period. The top section of the graph displays the Friedman statistics, revealing the p-value for the entire analysis and the Kendall coefficient of concordance along with 95% confidence intervals. These intervals help determine the effect size of the model. Brackets on the plot show Holm adjusted significant differences between week periods, determined using the pairwise test Durbin-Conover pairwise.

The performance during the two-week periods also exhibited variability between the two sites (Fig. S3). Notably, all Friedman tests yielded a low Kendall coefficient, indicating that although the samples differed significantly, there is likely another variable that could better explain this difference than the week of collection. The assessment of trap positions within the sites also revealed no significant differences (Fig. S4).

### Community composition

The composition of bee communities captured in traps of different color (all data combined) did not differ significantly (PERMANOVA: R^2^=0.035, p=0.011) (Fig. 5). While a significant p-value was obtained, the low R^2^ value indicates that the dissimilarity between color traps explains only a small portion of the variations in community composition, also the pairwise comparison showed not significant values. This is further supported by the proximity of centroids in the NMDS graph and the non-significant ANOSIM results (Fig. 5). However, the multilevel pattern analysis identified 14 species associated with a specific color trap (Table 2).

**Table 2.** Multilevel pattern analysis showing species that are strongly associated with a specific color trap (both sites combined).

| Trap color | Species | stat | p-value |
| --- | --- | --- | --- |
| Blue | *Andrena cineraria* (Linnaeus, 1758) | 0.238 | 0.0173 |
|  | *Nomada panzeri* (Lepeletier, 1841) | 0.238 | 0.0207 |
| Yellow | *Bombus humilis* (Illiger, 1806) | 0.343 | 0.0005 |
|  | *Bombus bohemicus* (Seidl, 1838) | 0.313 | 0.0017 |
|  | *Bombus barbutellus* (Kirby, 1802) | 0.243 | 0.0245 |
| Violet-white | *Bombus hypnorum* (Linnaeus, 1758) | 0.310 | 0.0013 |
|  | *Bombus jonellus* (Kirby, 1802) | 0.286 | 0.0045 |
|  | *Lasioglossum lativentre* (Schenk, 1853) | 0.216 | 0.0468 |
| White | *Bombus terrestris* (Linnaeus, 1758) | 0.334 | 0.0027 |
|  | *Bombus rupestris* (Fabricius, 1793) | 0.263 | 0.0108 |
|  | *Osmia bicornis* (Linnaeus, 1758) | 0.236 | 0.0272 |
| Yellow | *Andrena nitida* (Müller, 1776) | 0.303 | 0.0025 |
|  | *Andrena grvida* (Imhoff, 1832) | 0.250 | 0.0139 |
| Blue-yellow | *Andrena minutula* (Kirby, 1802) | 0.315 | 0.0025 |

**Fig. 5.**
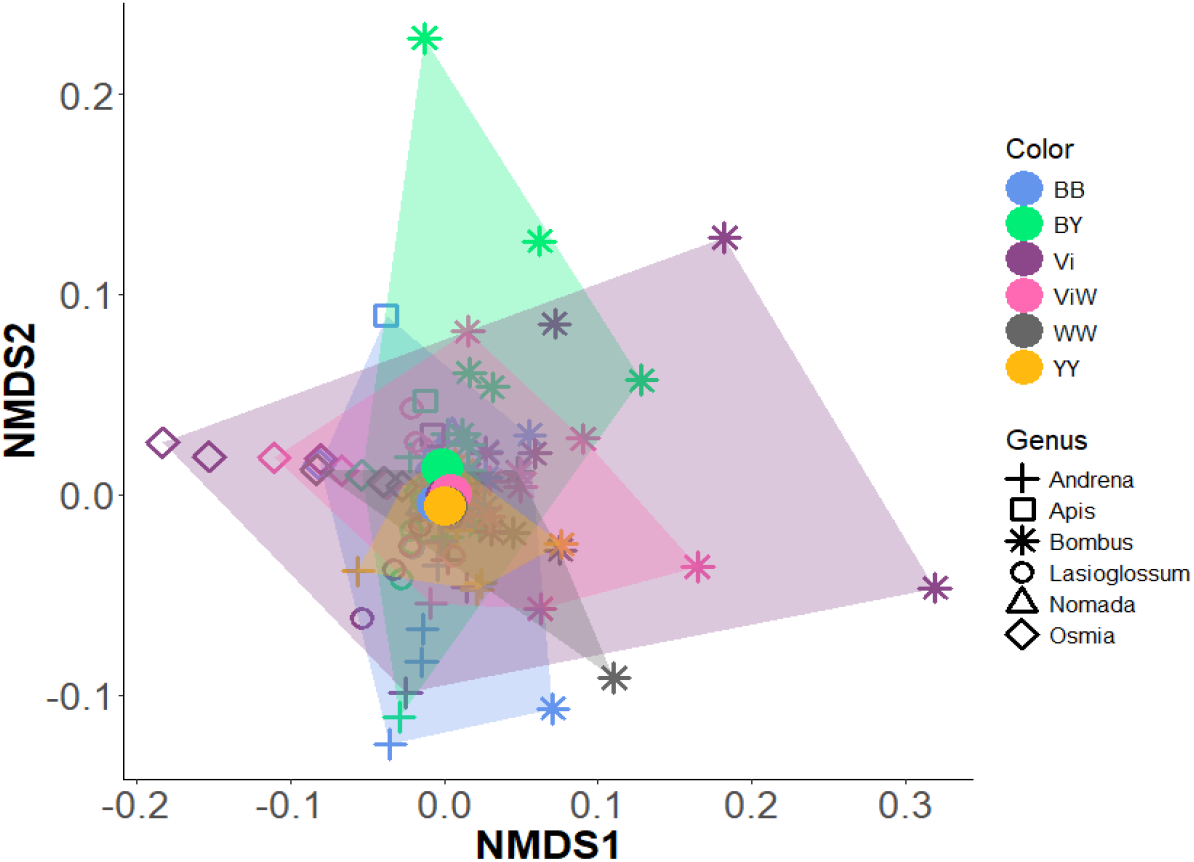
Two-dimensional non-metric multidimensional scaling ordination of bee communities collected with six different color trap types (sites H+HS combined), using Bray-Curtis dissimilarities (stress = 0.15). The color dots represent the different color traps, and the shapes represent the main genus of bees found in the traps.

Although the community composition pattern changed within each of the two sites, the lack of differences between color traps persisted (Fig. S5). The multilevel pattern analysis for the Haselberg habitat revealed nine associated species, while the Haselberg-slope habitat showed twelve species (Table S3). No significant differences in community composition were observed between different weeks on both sites and individually within each site.

Between the sites there was no significant dissimilarity in community composition (Fig. S6), but the multilevel pattern analysis identified eight associated species associated to a specific site (Table S4).

### Diversity

The diversity indices were compared across six different color traps from two habitats (H and HS). Blue-yellow had the highest observed number of species (41). However, violet exhibited the highest species richness (q=0) values, with 36 observed species, resulting in an estimated richness of 61 species. Despite this, confidence intervals across all traps overlapped, indicating no significant difference in species richness (Fig. 6a). Yellow had the highest Shannon diversity estimate values (which is sensible to common and rare species) with 39 estimated species (Table 3), yet yellow samples coverage was the lowest one. Yellow Shannon diversity was only significantly different to white. Yellow also had the highest Simpson diversity estimate value (which is sensible to common species) with 28 estimated species, and it was significantly different to white, blue and violet. Regardless, blue-yellow had the second higher q=1 and q=2 and higher sample coverage, meaning that the calculation is more reliable than with yellow (Table 3). Also, it was significantly higher than white, blue and violet (Fig. 6).

**Table 3.** Observed and estimated number of species (H and HS) according to species richness (q=0), Shannon diversity (q=1) and Simpson diversity (q=2) of the six different color trap combination. n is the total amount of individual observations; sample coverage is a is a metric used to measure the completeness of a sample.

| Trap color | $n$ | Sample coverage | Observed sp. | Estimated $q=0$ | Estimated $q=1$ | Estimated $q=2$ |
| --- | --- | --- | --- | --- | --- | --- |
| Blue | 119 | 0.88 | 33 | 55 | 24 | 13 |
| Blue-yellow | 141 | 0.90 | **41** | 53 | 35 | 25 |
| Violet | **159** | 0.90 | 36 | **61** | 24 | 13 |
| Violet-white | 125 | 0.90 | 33 | 47 | 26 | 17 |
| White | 74 | 0.79 | 27 | 59 | 23 | 10 |
| Yellow | 55 | 0.71 | 28 | 59 | **39** | **28** |

**Fig. 6.**
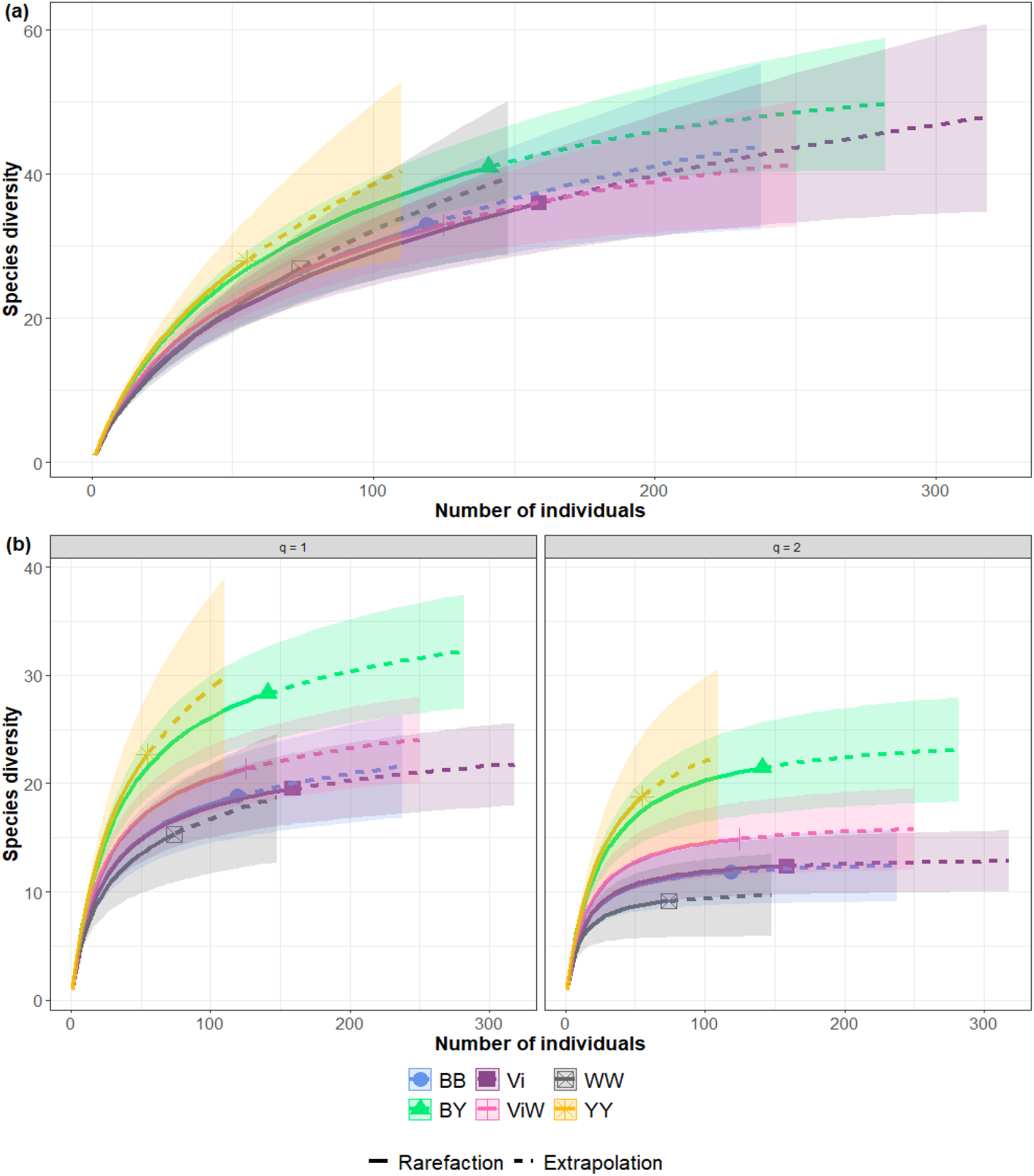
Sample-size-dependent Hill numbers. a) Species richness (q=0), (b) Shannon diversity (q=1) and Simpson diversity (q=2) rarefaction-estimation curves, with 95% confidence intervals (bootstrap 100 repetitions) of six color trap combinations at both habitats combined (H and HS). Trap colors: BB=Blue, BY=Blue-yellow, Vi=Violet, ViW=Violet-white, WW= White, YY= Yellow.

Analyzing bee diversity by habitat revealed substantial differences. At the Haselberg site (Fig. S7, Table S5), violet traps had the highest observed number of species (23) but white traps had most estimated species (36), significantly surpassing the richness of yellow traps. Shannon and Simpson diversity estimators were also highest in white traps, (26 and 17 species, respectively). However, because white had the lowest sample coverage, results from violet and violet-white traps are more reliable. At the Haselberg-slope site (Fig. S8, Table S5), blue-yellow traps had the highest observed number of species (32). Yellow had the highest species richness, Shannon and Simpson diversity estimators (127, 60 and 24, respectively), but also by far the lowest sample coverage. Comparing the two habitats, Haselberg-slope had more observed species and higher diversity values (Fig. S9, Table S6).

## Discussion

This study provides insightful observations on the color preferences of bees, suggesting that attractiveness of different colors is not only based in intrinsic bee color preferences but is likely to be also influenced by the habitat and their differing flower communities. Even in nearby habitats, distinct color preferences emerge.

Our findings indicate that violet and blue-yellow traps were the most successful, with higher abundance and observed species richness across both habitats. Particularly, violet proved to be the most successful color in the Haselberg site, while blue-yellow dominated in the Haselberg-slope site. Despite some studies deeming violet traps redundant (Acharya et al. 2022), we decided to include in our study. We believed that violet, being a naturally occurring color in nature, would find success in a habitat with plenty of violet flower species. In contrast to previous studies, our violet traps achieved notable success, prompting the consideration of various factors such as color hue and habitat specificity in trap design and deployment strategies. We emphasize that our study has used a ‘true’ violet - ultramarine violet - i.e. not one of the commonly used mixtures of red and blue pigments that appears violet in a human’s eye be not in the eye of an insect. The difference between ‘true’ and ‘false’ violet might also explain why violet has not been very successful in previous studies.

While blue is a commonly used color for bee collection (Gibbs et al. 2017; Kimoto et al. 2012; Stephen and Rao 2007; Turley et al. 2022), our study aligns with others showing variations of color effectiveness (Abrahamczyk et al. 2010; Buffington et al. 2021; Gollan et al. 2011; Grundel et al. 2011; Hall 2016; Saunders and Luck 2013; Sircom et al. 2018; Vrdoljak and Samways 2012), and this variation can be explained by the diverse habitats the studies took place in. The habitat factor showed to be of crucial importance in this study (Fajemisin et al. 2023; Saunders and Luck 2013; Westerberg et al. 2021). Bee floral preferences are linked to dominant flora, phenology, and shape of an area, with different flower colors gaining prominence in different habitats (Abrahamczyk et al. 2010; Fajemisin et al. 2023; Freitas-Moreira et al. 2016; Saunders and Luck 2013). Although bees generally favor UV, blue, and green spectra (Chen et al. 2020; Gumbert 2000; Ostroverkhova 2018; Simonds and Plowright 2004), learned color preferences from floral resources can override this predisposition to a certain extent. (Freitas-Moreira et al. 2016; Gumbert 2000). Given that bees are often generalist feeders, successfully identifying flowers that offer the most significant rewards is crucial for optimizing their foraging time. Consequently, color emerges as one of the primary factors that bees learn to discriminate (Chittka and Menzel 1992; Chittka and Thomson 2001).

Another factor is the difference in hue and reflectance of the colors used across studies (Acharya et al. 2022; Krčmar et al. 2014; Vrdoljak and Samways 2012), this difference could account for some variations in attractiveness for bees. A clear example of the intricate relationship between flowers and color preference is seen in oligolectic bee species, especially those that feed on similarly colored flowers within a plant genus. In such cases, it is highly likely that these bees will choose a color trap resembling the hues of the flowers they predominantly feed on (Heneberg and Bogusch 2014; Leong and Thorp 2001; Pickering and Stock 2003). In our study, we observed several violet-colored flower species consistently appearing during most of the six-month period, emphasizing the significance of color preferences in bee foraging behavior.

The blue-yellow trap stood out as a significant finding in our study, ranking second in overall bee abundance and the first in HS, also capturing the highest number of bee species. This success prompts further investigation and highlights the importance of contrast in attracting bees to flowers, and the blue-yellow trap efficacy may be attributed to its ability to provide a high-contrast target for bees. Contrast is vital for correct flower identification, and bee-pollinated flowers typically share a characteristic: UV-absorbing centers and UV-reflecting peripheries (Koski and Ashman 2014; Papiorek et al. 2016; van der Kooi et al. 2019). In contrast, flowers mainly pollinated by birds lack color contrast and spectral purity (de Camargo et al. 2019; Papiorek et al. 2016). Furthermore, using two colors in one trap could potentially overcome habitat-specific biases by presenting a higher number of colors, thus attracting a more comprehensive community of bees. This approach may contribute to increased collection efficiency with fewer traps.

We observed no significant differences in community composition among the traps, a result that aligns with certain previous studies (Buffington et al. 2021; Hall 2016; Toler et al. 2005) while differing from others that reported substantial disparities (Freitas-Moreira et al. 2016; Gollan et al. 2011; Hall 2018; Joshi et al. 2015). Despite this, we identified specific bee species exclusively collected in traps of certain colors. Interestingly, the color preference of these species did not strongly correlate with the described bee-flower associations in Germany (Westrich 2019).

Noteworthy was the observation in *Andrena*, primarily captured in blue, yellow, and blue-yellow traps. For instance, *A. bicolor*, one of the most abundant species, exhibits a highly generalist feeding behavior, foraging from around 15 plant families (Westrich 2019). This generalist behavior provides a rationale for its attraction to blue and yellow. Blue-yellow emerged as a versatile option, potentially serving as a single trap effectively capturing a diverse array of species, offering both independent color attractiveness and a combined effect. It is essential to highlight that blue-yellow represented nearly all Andrena species collected in the study.

It is important to note that while our color traps successfully captured a substantial number of bee species in the area, there were additional species present that our traps did not collect. While color is a primary factor influencing flower discrimination for bees due to its visibility from a distance, other characteristics such as plant height, flower shape, size, and odor also play crucial roles (Chittka and Menzel 1992). For instance, carpenter bees (*Xylocopa*), known to forage during twilight, rely on olfactory cues to identify their preferred flowers (Araújo et al. 2024). This highlights the importance of considering multiple factors beyond color alone in bee foraging behavior.

Bee abundance fluctuations during the collection period did not correlate significantly with weather conditions (temperature, humidity and precipitation) but were more influenced by phenological changes. Two peaks in abundance were observed, corresponding with the activity of certain bee species. One peak was in late March and early April, coinciding with the appearance of *O. bicolor* and some hibernating *Bombus* species. Another peak was in mid-July, aligning with the emergence of young workers and males from various *Bombus* species (Westrich 2019). The dominance of the genus *Bombus* and the species *O. bicolor* as the most abundant ones suggests that their phenology significantly contributed to the changes in bee abundance throughout the collection period. Bees have been found to be more influenced by the quantity and diversity of flowers in their habitat than by temperature, unlike other insect groups (Abrahamczyk et al. 2011; Fajemisin et al. 2023).

Our study underscores the importance of considering habitat-specific dynamics in trap design and the potential of violet and blue-yellow traps. Future research should explore using tailored color combinations and additional trapping methods to achieve a comprehensive representation of bee communities

## Conclusions

This study demonstrates that bee color preferences in vane traps are influenced not only by intrinsic visual biases but also by habitat-specific factors. Our study highlights the influence of both color contrast and habitat-specific factors on bee trapping effectiveness. Violet and blue-yellow traps captured the highest bee abundance and diversity, with blue-yellow emerging as the most versatile option, likely due to its high contrast and ability to appeal to a broader range of bee species. The observed differences in community composition between two nearby habitats underscore the impact of local environmental factors on bee behavior. This suggests that floral availability, landscape structure, and local environmental factors play a role in shaping bee responses to color cues. These findings emphasize the need for tailored trap designs that account for both color contrast and habitat dynamics. Future research should explore additional color combinations and alternative trapping methods to improve species representation in biodiversity surveys.

## Data Availability

Data from this work and supporting material is available at: https://osf.io/3ytgj/?view_only=b0a393c0f986445e9ef04ba23aa2b267

## Acknowledgements

Special appreciation goes to Frank Creutzburg for sparing some of his time to assist in the process of bee identification, contributing to the research. Daniel Veit thankfully helped with the construction of the vane traps, and the UNB Jena (Frank Hünefeld) issued the required collection permits.

## Author information

### Authors and Affiliations

Institut für Zoologie und Evolutionsforschung mit Phyletischem Museum, Friedrich-Schiller-Universität Jena, Jena, Germany

Elizabeth Rentería and Gunnar Brehm

### Contributions

The study was conceived and designed by Gunnar Brehm and Elizabeth Rentería. The experiment was set up in the field by GB and ER. Data collection in the field was carried out by ER. Data analyses was performed by ER; the spectral analysis was performed by GB. The first draft of the manuscript was written by ER, and it was reviewed, edited, and approved by both authors.

## Ethics declarations

### Ethics

Efficient collection and identification required the killing of bee (and other insect) specimens. The approach is justified by the gain of important data. i.e. species lists that are otherwise not available, and can help with management decisions in the conserved area. Bee specimens are permanently deposited in the Phyletisches Museum and are available for further study.

